# Assembly and Cation-selectivity Mechanisms of Neuronal Gap Junction Channel Connexin 36 Elucidated by Cryo-EM

**DOI:** 10.1101/2023.04.24.537958

**Authors:** Wenxuan Mao, Shanshuang Chen

## Abstract

Electrical synapses are essential components of neural circuits. Dysfunction of electrical synapses can lead to obstacles in learning and memory. Neural signal transduction across electrical synapses is primarily mediated by a gap junction channel, Connexin 36 (Cx36), the lack of which causes impaired electrical couplings in cortical interneurons and thalamic reticular nucleus (TRN) neurons. Unique characteristics of Cx36 gap junction channel include its insensitivity in transjunctional membrane potential, as well as its preference for homomeric assembly, prone to exclude other paralogous connexins from co-assembly. However, the structural basis underlying Cx36 function and assembly remains elusive. Here, we report the cryo-EM structure of human Cx36 at 2.67 Å resolution and identified critical residues underpinning its obligatory homomeric assembly. In particular, we found non-canonical electrostatic interactions between protomers from opposing hemichannels and a steric complementary site between adjacent protomers within a hemichannel, which together offer a structural explanation for the assembly specificity in homomeric and heteromeric gap junction channels. Moreover, the narrowest restriction along the channel axis overlaps with an acidic patch, where Glu43, Asp47 and Asp48 may contribute to cation-selectivity. Also, the amino-terminal helix reported to be responsible for sensing membrane potential in other connexins was disordered in our Cx36 structure, in line with its channel activity independent of membrane potential. Together, this work elucidated the assembly mechanisms of the electrical synaptic gap junction channel, and offered possible explanations for sustained Cx36 activity upon membrane depolarization, which allows efficient action potential propagation across electrical synapses.

## Introduction

Synapses are the key sites for neurotransmission in the nervous system. In addition to the well-recognized chemical synapses, emerging evidences suggest that electrical synapses are also indispensable for a variety of neural activities including learning and memory. Electrical synapses are often found in inhibitory interneurons in diverse regions of the mammalian central nervous system (CNS) [1], contributing not only to the development of neural circuits [2-4], but also the transduction of membrane depolarization by directly allowing the permeation of ions [5]. Studies of olfactory bulb [6], cerebral cortex [1, 7], thalamic reticular nucleus [8] and inferior olivary nucleus [9, 10] also revealed a special role of electrical synapses as the mediator of synchronization between connected neurons. In contrast to chemical synapses, electrical synapses lack distinct morphological specifications under electron microscopes such as presynaptic active zones and post-synaptic densities, and thus are more difficult to identify in brain slides [11-14]. The cleft of electrical synapses is usually smaller than chemical synapses, such that gap junction channels can span the membranes of two adjacent neurons to directly mediate electrical signal transduction [15]. Gap junction channels are homomeric or heteromeric dodecamers, formed by two hexameric hemichannels (also known as connexons) docked in dihedral symmetry, each localized in the membrane of adjacent cells [16]. Connexin gene family is highly diverse, and many connexins have been found in electrical synapses. Cx36, the most abundant subtype, is thought to function in sole electrical synapses, whereas other subtypes including Cx45 [17], Cx57 [18] and Cx50 [19] are present in electrical-chemical “mixed” synapses [20]. Genetic ablation of Cx36 leads to near complete loss of electrical couplings in cortical interneurons and thalamic reticular nucleus (TRN) neurons [21, 22], and functional deficits such as visual [23], motor [24], and memory impairments [25] have also been observed. Similarly, Cx36 is also responsible for the electrical coupling between pancreatic β-cells [26]. The molecular mechanisms of Cx36 remain to be investigated, and the structural understanding of electrical synapses.

In addition to the neuronal Cx36, astrocytes and oligodendrocytes are found to express other connexins such as Cx26 [27], Cx32 [28, 29], Cx43 and Cx47 [30], which could form heteromeric assemblies to conduct ions [5], small peptides, secondary messengers or metabolites [31-34]. However, Cx36 is prone to form homomeric assemblies between neurons in hippocampus, olfactory bulb, retina and spinal cord neurons, and possibly other regions in the CNS [35-38], consistent with the notion that Cx36 is dedicated for electrical synapses. Moreover, Cx36 features low conductivity and transjunctional voltage insensitivity, distinct from other connexins [39], also in line with its primary role in faithfully and efficiently propagating electrical pulses across the electrical synapses. Given the high sequence similarities between Cx36 and other connexins, it thus requires structural determination to elucidate the mechanisms by which Cx36 does not co-assembly with other paralogous connexins, and how its conductivity and voltage sensitivity might be rendered. In this study, we report the structure of Cx36 homododecamer determined by cryo-electron microscopy (cryo-EM), which offers reliable mechanistic insights for Cx36 assembly and may shed light on how cation selectivity and voltage insensibility of neuronal connexins are achieved.

## Methods

### Plasmid construct

The full-length cDNA of human Cx36 (National Center for Biotechnology Information reference sequence: NP_065711.1) was cloned into pBacMam expression vector in fusion with a C-terminal eGFP and a Strep II-tag for subsequent over-expression and purification experiments [40].

### Baculovirus production

Baculovirus used for overexpression of Cx36 in Expi293F cells was obtained using the standard Bac-to-Bac system. In brief, the aforementioned pBacMam plasmid construct was transformed into DH10Bac cells. Recombinant bacmid clones containing Cx36 overexpression cassette were extracted from white colonies selected on LB agar plates supplemented with 50 μg/ml kanamycin, 7 μg/ml gentamicin, 10 μg/ml tetracycline, 100 μg/ml Bluo-Gal and 40 μg/ml IPTG. After incubation at 37 °C for 48 hours, a single white colony was inoculated into 5 ml LB medium containing 50 μg/ml kanamycin, 7 μg/ml gentamicin and 10 μg/ml tetracycline. Cells grown overnight were centrifuged at 4000 rpm (Allegra X-12R, Beckman Coulter) for 10 min at 4 °C. The cell pellet was resuspended in 200 μl P1 buffer (QIAGEN Miniprep Kit), followed by addition of 200 μl P2 buffer for cell lysis. After the suspension became clear, 200 μl N3 buffer was added, and the mixture was centrifuged at 14000 rpm (Centrifuge 5418R, Eppendorf) for 10 min. Supernatant was mixed with 1 ml isopropanol and incubated in -20 °C for more than 1 hour to precipitate bacmid DNA. After centrifugation at 14000 rpm for 10 min, pellet was washed by 70% ethanol and then resuspended in 50 μl ddH_2_O. To produce P1 Baculovirus, 1 μg bacmid dissolved in ddH_2_O was mixed with 8 ul of Cellfectin II (Thermo Fisher Scientific) before added to Sf9 cells seeded at 0.9×10^6^ cells/ml. P1 virus was harvested by filtering the supernatant media through a 0.2 μm filter. P2 virus was subsequently generated by adding P1 virus to Sf9 cells at a 1:2000 (v/v) ratio when Sf9 cells reached 1.2×10^6^ cells/ml. P2 virus was harvested by filtration after 4 days. For preservation, 2% (v/v) fetal bovine serum (FBS) was added to P1 and P2 virus before filtration.

### Cx36 over-expression

For large-scale expression, Expi293F cells cultured in suspension at 37 °C, 5% (v/v) CO_2_ with gentle shaking (110 round per minute (rpm)) were infected by P2 virus at a final concentration of 8% (v/v) when cell Expi293F density reached 2.5×10^6^ cells/ml. Sodium butyrate was supplemented to a final concentration of 10 mM 12 hours after infection and cell culture were thereafter transferred to a different incubator set at 30 °C, 5% (v/v) and 110 rpm. Cells were cultured for another 48 hours before harvest by centrifugation at 4,000 rpm for 15 min (JLA-8.1 rotor, Beckman Coulter). Cells were washed twice with phosphate buffered saline (PBS) and stored at -80 °C until use.

### Cx36 purification

Thawed cell pellets were resuspended and homogenized in lysis buffer containing 25 mM HEPES pH7.4, 500 mM NaCl, 1% (m/v) DDM and 0.2% (m/v) CHS for 2 hours at 4 °C. After ultra-centrifugation at 40,000 rpm for 1 hour in (Ti45 rotor, Beckman Coulter), the supernatant was incubated with Streptactin-XT resin equilibrated with Buffer A (25 mM HEPES pH7.4, 150 mM NaCl, 1 mM DDM and 0.2 mM CHS) for 30 min at 4 °C with gentle rocking. Thereafter, the resin was collected into a column and rinsed by Buffer A for 3 column volumes. Purified Cx36 was eluted with Buffer A supplement with 150 mM biotin. Eluted protein was subjected to a desalting column (PD-10 Columns, GE Healthcare) to remove biotin, and subsequently digested with 3C protease at a ratio of 1:1 (m/m) at 4 °C overnight. Incompletely digested protein and free GFP were thereafter removed by passing through regenerated Streptactin-XT column. The flow-through was concentrated using 100 kDa cut-off concentrator (Millipore) before loaded onto an SEC column (Superose 6 Increase 10/300 GL) pre-equilibrated with SEC running buffer containing 25 mM HEPES, 150 mM NaCl and 0.03% GDN. Peak fractions containing Cx36 were pooled and concentrated to ∼4.5 mg/ml for cryo-EM sample preparation.

### Cryo-EM sample preparation

Right before use, Cx36 sample was centrifuged at 14000 rpm (Centiufuge 5418R, EPPENDORF) for 10 min at 4 °C to remove any debris. Grids (Quantifoil Cu R1.2/1.3, 300 mesh) were glow discharged at 15 mA for 30 s using PELCO easiGlow system. When the vitrobot (Vitrobot Mark IV, Thermo Fisher Scientific) chamber is equilibrated to 8 °C and 100% humidity, 5 μl sample was applied to grids. After a 10 s incubation, grids were blotted for 3.5 s and subsequently plunged into liquid ethane cooled by liquid nitrogen. Frozen cryo-EM samples were stored in liquid nitrogen until use.

### Cryo-EM data acquisition

Grids were first screened in a 200 kV electron-microscope (Talos Arctica, Thermo Fisher Scientific) of 200 kV acceleration voltage, equipped with a direct electron detector (Falcon IV, Thermo Fisher Scientific). Samples of desired ice thickness were used for data collection in a 300 kV electron-microscope (Titan Krios G3i, Thermo Fisher Scientific) equipped with a cutting-edge direct electron detector (K3, Gatan). Movies were collected at a nominal 81,000× magnification in super-resolution mode where the magnified physical pixel size was 1.1 Å and a nominal defocus value ranging from 1.5 to 2.5 μm was applied. A total dose 50 e^-^/Å^2^ was evenly fractionated into 32 frames, and a dataset composed of 1817 micrographs was collected.

### Image processing

Movie stacks were corrected for beam-induced drift using MotionCor2 [41] and contrast transfer function (CTF) parameters per micrograph were estimated in GCTF [42]. Particle picking was carried out using Gautomatch (https://www2.mrc-lmb.cam.ac.uk/download/gautomatch-056/) with reference images generated by projection of reported Cx26 reconstruction in various directions. An initial set of 883052 particles was extracted with a box size of 280 pixels in Relion-4.0 [43]. After several rounds of 2D classification performed to remove ice and contaminants, 316019 good particles were selected for subsequent 3D classification using C1 or D6 symmetry. The reference map used for 3D classification is from Ab-initio reconstruction in cryoSPARC-V3.3.2 [44]. The class with reasonable connexin dodecamer density features containing 117469 particles was used for subsequent 3D refinement with D6 symmetry in Relion. To improve the resolution, CTF refinement and Bayesian polishing were performed and yielded a final map at 2.77 Å resolution determined by gold standard Fourier shell correlation (FSC). This set of particles was also applied to cryoSPARC for an additional round of homogeneous refinement, generating a reconstruction at 2.67 Å.

### Model building

The initial model of human Cx36 was generated by homologous modelling from a Cx26 structure (PDB code 7QET). This model was then fitted into the density map as a rigid body using UCSF chimera. Manual adjustments were carried in COOT [45], where clear density feature could guide unambiguous structural modelling. The model was then subjected to real-space refinement in Phenix [46]. After several rounds of modelling and refinement, the final model was validated by MolProbity [47]. Statistics are described in Extended Data Table 1.

### Figure preparation

All structural figures were prepared using Pymol and UCSF chimera.

## Results

### Structure determination of CX36 in detergent

To understand the function and assembly of Cx36 at electrical synapses, we carried out single particle cryo-EM studies of the full-length human Cx36, and determined its three-dimension structure to 2.67 Å (Fig. 1a). Initially extracted in detergent micelles containing N-Dodecyl-β-D-maltopyranoside (DDM) and cholesterol hemisuccinate (CHS), affinity chromatography-purified Cx36 was cleaved to remove eGFP and the affinity tag, and then subjected to detergent exchange into glycodiosgenin (GDN) by size-exclusive chromatography (SEC), where it eluted at a volume consistent with dodecameric assembly (Extended Data Figs. 1a and 1b). Resulted peak fraction was thereafter used for cryo-EM data collection. Electron micrographs are of decent contrast, and characteristic “dumbbell”-shaped gap junction channels can be unambiguously recognized as monodisperse single particles (Extended Data Fig. 1c). Two-dimensional class averages of picked particles showed evenly distributed projection angles, and confirmed the dodecameric assembly of Cx36 (Extended Data Fig. 1d). Surprisingly, besides most class averages obeying a six-fold symmetry about the central axis, a few classes of “top-down” projections are seven-fold related, possibly with pseudosymmetry (Extended Data Fig. 1d). We reasoned that these minor classes could represent an artificial assembly resulted from detergent extraction, and thus excluded these particles from further analysis. With imposed D6 symmetry, we obtained a final three-dimension reconstruction at an overall resolution of 2.67 Å (Extended Data Figs. 2a, 2b, 2c and 2d). The outstanding map quality allowed accurate model building of amino acids from 17 to 103 and 186 to 285, whereas the aminoterminal region (Residues 1-16), the cytoplastic loop (Residues 104-185) and the carboxyl-terminus (Residues 286-321) were disordered (Fig. 1b and Extended Data Fig 2e).

**Fig. 1.**
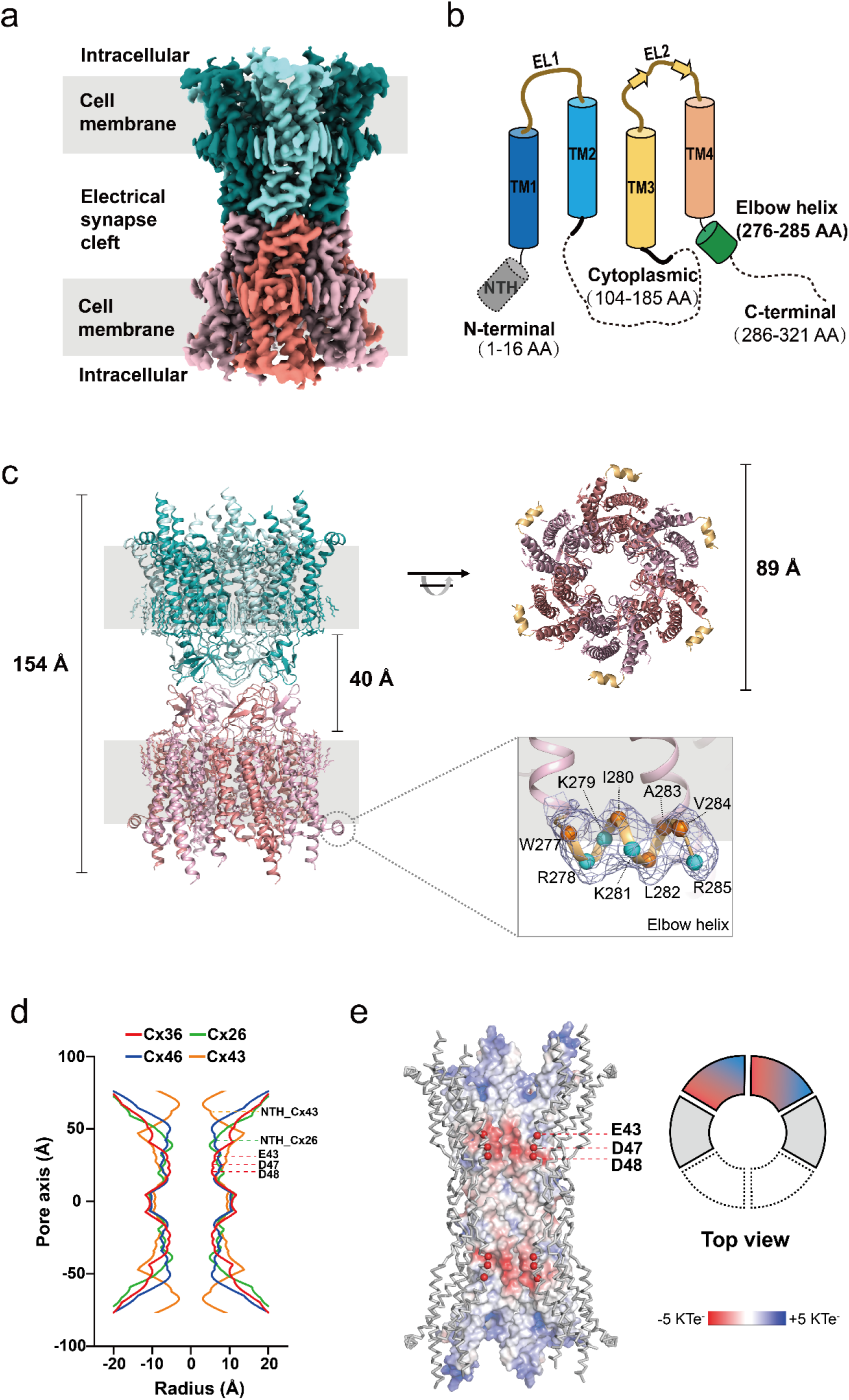
Cryo-EM structure of gap junction protein Cx36. **a**, Overall cryo-EM map of the Cx36 shown in front view. Adjacent protomers in upper hemichannel are colored in teal and pale cyan, whereas the lower hemichannel are colored in salmon and light pink. Region corresponding to the cell membrane is indicated in gray. **b**, Schematic representation of the topology of a Cx36 protomer. Cylinders and arrows indicate α-helices and β-strands, respectively. Structurally disordered regions are indicated by dashed lines. **c**, Atomic model of Cx36 dodecamer displayed in ribbon representation, in front view (left) and top-down view (upper right). An amphipathic elbow helix in carboxyl terminal is shown in lower right corner, where the Cα atoms of hydrophobic and polar residues are colored in orange and pale cyan, respectively. Map density for the elbow helix is shown as blue mesh. The gray background represents the cell membrane. **d**, Solvent-accessible pore radius profiles calculated for Cx36 (red, PDB code 8IYG), Cx26 (green, PDB code 2ZW3), Cx46 (blue, PDB code 6MHQ) and Cx43 (orange, PDB code 7Z1T) using the HOLE program is superimposed for comparison. The positions corresponding to Glu43, Asp47, and Asp48 of Cx36 are indicated with red dashed lines, while the positions of NTH of Cx26 and Cx43 are marked with green and orange dashed lines, respectively. **e**, Electrostatic potential surface calculated using Adaptive Poisson-Boltzmann Solver is shown for the two Cx36 protomers in the back, whereas the two side protomers are shown in gray Cα traces and the two front protomers are hided for clarity. The Cα atoms of Glu43, Asp47, and Asp48 are represented by red spheres.

The overall architecture of Cx36 is largely conserved among known gap junction channel structures. The barrel-shaped dodecameric channel consists of two hexameric hemichannels that docked with each other on the extracellular side. In comparison, the dimensions of Cx36 (δ-subfamily connexin) are 154 Å by 89 Å by 89 Å (Fig. 1c), close to those of the Cx26 (β-subfamily connexin) and Cx46 (α-subfamily connexin), with pairwise Cα root mean squared deviations (r.m.s.d.) of 1.34 Å and 1.25 Å, respectively. The channel pore profile analyzed by HOLE program [48] is also similar, with acidic residues Glu43, Asp47 and Asp48 lining up the pore (Fig. 1d). Despite the high local resolution and good side chain density of most neighboring residues, there is only reliable density for the main chain of these acidic residues, suggesting side chain flexibility at least in part due to electrostatic repulsion (Fig. 1e). Nonetheless, these D6-symmetry related pore-lining acidic patches, one in each protomer, would repulse anions from entering, thus rendering Cx36 gap junction channel highly selective for cations. They may also contribute to the blockage by multiple bivalent cations. Consistently, Asp47 has been report to render Cx36 gap junction sensitive to magnesium [49, 50]; however, we failed to observe any putative magnesium density coordinated by the carboxyl groups of Asp47 side chains, or in a different reconstruction at 2.9 Å resolution using sample prepared in the presence of 5 mM magnesium (data not shown).

On the other hand, our structure revealed a few minor but distinct features. One is that the first 16 residues are disordered in our structure. By contrast, this region in other connexins forms a so-called amino-terminal helix (NTH) located in the cytoplasmic entrance of the gap junction channel, thought to be responsible for channel regulation by membrane potential [51, 52]. The equivalent NTH binding sites in Cx36 are occupied by putative CHS, lipids or detergent molecules, (Extended Data Fig.4), in line with the observation that Cx36 operates with little dependence on membrane potential, although we could not rule out the possibility that the NTHs are removed from their consensus sites during detergent extraction. The other distinct structural feature of Cx36 is a short helix at the cytoplasmic end of the fourth transmembrane helix (TM4) and parallel to the membrane plane, which we deemed elbow helix. The elbow helix appears to be amphipathic, with hydrophobic Trp277, Ile280, Ala283 and Val284 located on the membrane side of the helix and hydrophilic Arg278, Lys279, Lys281 and Arg285 located on the cytoplasmic side (Fig. 1c). We speculate that these amphipathic elbow helices function to stabilize Cx36 in the membrane. Indeed, Arg215 in human Cx32, equivalent to Arg278 in Cx36, abolishes gap junction channel function when mutated to tryptophan, suggesting importance of this position [53].

### Distinct hemichannel docking mechanism of Cx36

Gap junction channel formation requires proper docking of two hemichannels, each from an adjacent cell. The extracellular loop (EL) regions of connexin hemichannels usually protrude beyond the cytoplasmic membrane of the cell by approximately 20 Å [54], thus the cleft of Cx36 containing electrical synapse can be as narrow as 4 nm, much smaller than that of a typical chemical synapse. Reported functional studies have not only characterized the gating and regulatory properties of many gap junction channels, but also accessed functional channel formation by homotypical or heterotypical assembly. The first structural insight into connexin assembly mechanism was obtained using Cx26 homododecamer [55], which revealed interactions between extracellular loops that governs hemichannel docking, and together with subsequently determined connexin structures, began to uncover docking mechanisms by which homotypical or heterotypical assembly is specified. Interactions between the opposing first extracellular loops (EL1) are primarily hydrogen bonds conserved in known connexin structures including Cx46/50 [56], Cx43 [57], the recently reported Cx36 [58], and Cx36 in this study. In our structure, these hydrogen bonds were formed by Asn56/Leu58 and their opposing partners in a head-to-toe fashion, and between dihedral symmetry-related Gln59 from a different protomer pair (Figs. 2a and 2b), highly consistent with previously reported Cx26. In comparison, the residues involved in hydrogen bonds in EL1 of Cx46/50 and Cx43 are different, yet the pattern that EL1 from each protomer simultaneously interacts with two neighboring protomers form the opposing hemichannel is conserved (Figs. 2c and 2d). Such pattern ensures a hydrogen bond network that strengthens not only the interactions between hemichannels, but those between protomers within a hemichannel as well.

**Fig. 2.**
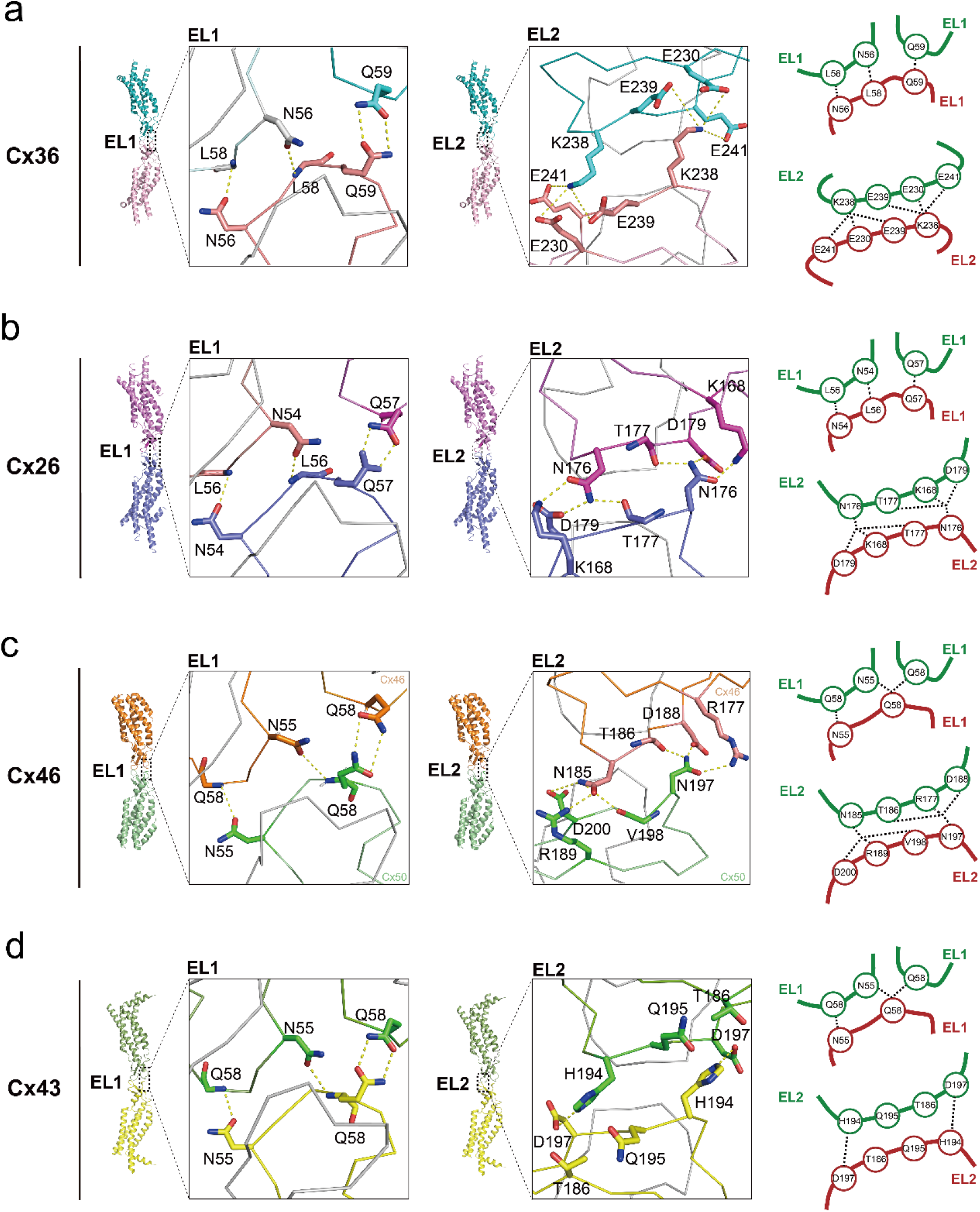
Hemichannel docking interface in Cx36, Cx26, Cx46 and Cx43. **a**, Zoom-in views of EL1-EL1 (left) and EL2-EL2 (middle) docking interface in Cx36. The key residues that mediate interactions at the interface are shown as sticks. Schematic representation of EL1-EL1 interaction (upper right) and EL2-EL2 (lower right) are shown for simplicity. **b, c, d**, Similar views of the docking interfaces and schematic representation for EL1-EL1 and EL2-EL2 interactions are shown for Cx26 (b), Cx46(c) and Cx43(d).

Unlike the EL1 interactions, the other extracellular loop EL2 interactions are constricted within opposing protomer pairs, thus likely contributing little to stabilization of adjacent protomers. Surprisingly, EL2 of Cx36 forms distinct interactions to stabilize the gap junction channel. Electrostatic interactions were observed where the positively charged Lys238 side chain is wedged in a negative patch composed of Glu230/Glu239/Glu241 from the opposing EL2, like a baseball in the glove, where the baseball is represented by Lys238 and the glove is formed by Glu230/Glu239/Glu241 (Fig. 2a). In sharp contrast, EL2 interactions in other connexin paralogs are largely mediated by hydrogen-bonds with the “baseball” position occupied by a neutral residue instead (Figs. 2b, 2c and 2d). Cx36 residues involved in the electrostatic interactions are poorly conserved, suggesting the uniqueness of the interaction and that Cx36 might exclusively form homotypical gap junction channels. Although residues contributing to EL2 interactions are highly diverse among connexins, their relative positions are perfectly consistent in the sequence alignment (Extended Data Fig. 3). Therefore, the “baseball in the glove” fashion of EL2 interactions can still be shared by all known connexins. To further quantify EL2 interactions, we measured the Cα distances from Lys238 to each interacting glutamate, resulting in 7.6 Å, 8.6 Å and 10.7 Å with Glu230, Glu239 and Glu241, respectively. These equivalent distances are 6.3 Å, 7.9 Å and 9.0 Å in Cx26, 5.7 Å, 7.3 Å and 8.7 Å in Cx46 and 6.3 Å, 8.2 Å and 9.5 Å in Cx43. These measurements revealed that the EL2 main chains between opposing protomers are slightly further than in other connexins, further reinforcing the distinction of EL2 interactions in Cx36. To validate the significance of above difference in distances, we compared the Cα distances between the conserved glutamines that form hydrogen bonds in EL1 across structural available connexins, and found they are near identical (6.5 Å in Cx36, 6.3 Å in Cx26, 6.6 Å in Cx46 and 6.6 Å in Cx43), thus ruling out any systematic error. In conclusion, the quantification shows that EL2 interactions in Cx36 are evolutionarily divergent in comparison with other connexins. Given the fact that the Lys238, Glu230, Glu239 and Glu241 are not seen in equivalent positions in any other connexins, it is likely the unique EL2 of Cx36 precludes heterotypical hemichannel docking.

### Hemichannel assembly mechanisms of Cx36

Assuming the unique EL2 structure of Cx36 prevents heterotypical interaction with a different connexin protomer at the opposing position, if Cx36 co-assembles with other connexin to form any heteromeric hemichannel, it would be a rare coincidence such a heteromeric hemichannel could dock with a matching hemichannel counterpart to give rise to a heterotypical and heteromeric gap junction channel. Prevention of Cx36-containing heteromeric hemichannel is thus necessary to ensure efficient utilization of Cx36 protomers, and can be achieved by transcription regulation. However, reported single-cell transcriptome showed that not all Cx36 positive neurons express Cx36 as the sole connexin, and that co-expression of Cx36 and other connexins is indeed detected [59]. We thus hypothesize that there must be an alternative mechanism for hemichannel assembly specificity intrinsic to connexins. To test, we examined the protomer interface within homomeric hemichannels of Cx36 and Cx26. We noticed that Arg233 in EL2 of Cx36 protrude its long and charged side chain towards Gly227 of the neighboring protomer, whereas such protrusion is reversed in Cx26, that is, the positions of bulky and small side chain residues are swapped, with Ala171 and Arg165 replacing aforementioned Arg233 and Gly227, respectively (Fig. 3a). We thus term these two spatially opposite modes of stereo complementation “short-R” for Cx36 and “R-short” for Cx26. Superposition of Cx36 and Cx26 hemichannels using a given protomer from each structure as a reference resulted in a Cα r.m.s.d. of 1.34 Å and thereafter a modeled Cx36/Cx26 heteromeric interface. We observed that the Arg233 from Cx36 clashes with Arg165 from Cx26, thereby destabilizing the modeled heteromeric interface (Fig. 3a). In summary, we hypothesize a stereo complementation mechanism that contributes to hemichannel assembly specificity.

**Fig. 3.**
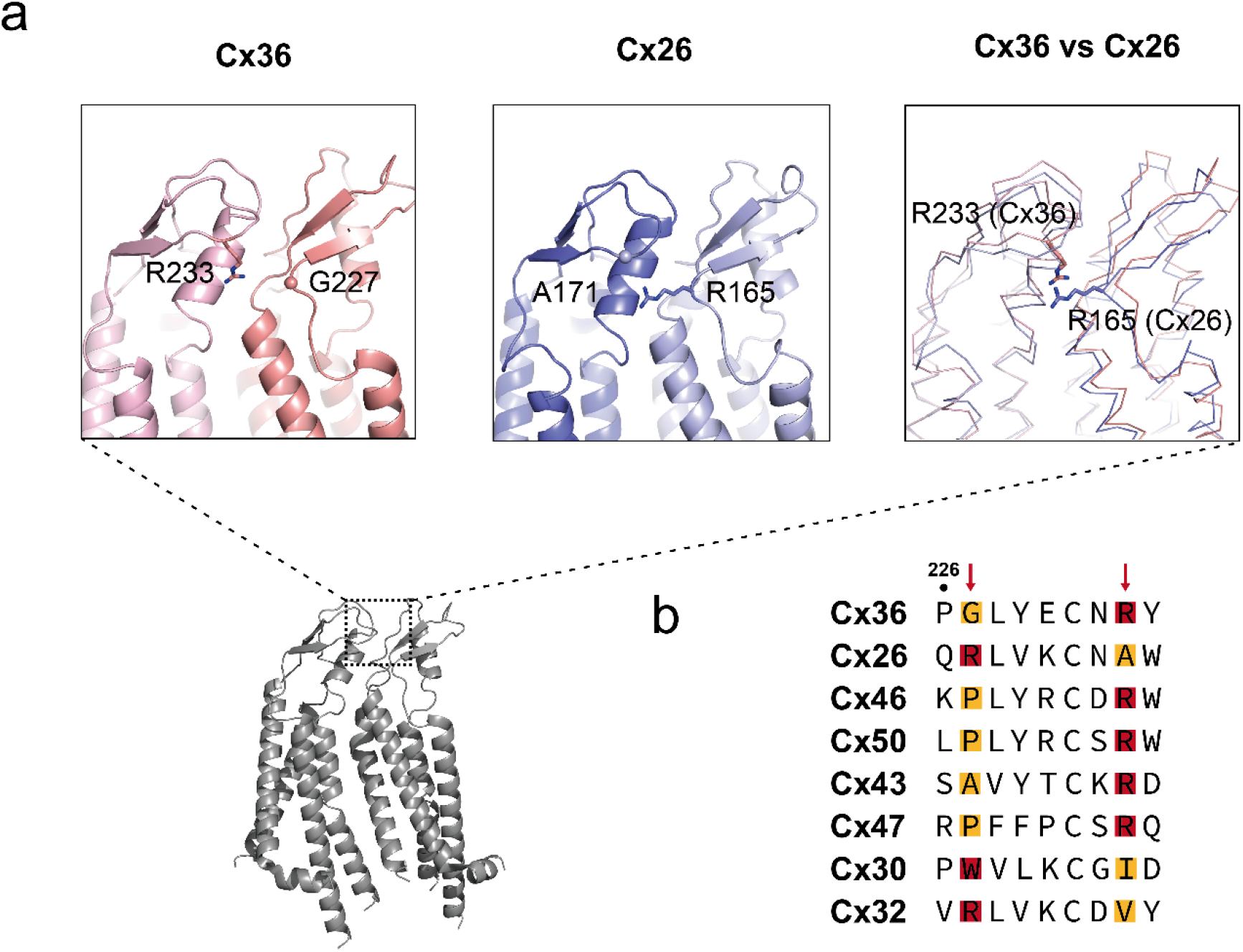
The docking interface between adjacent protomers. **a**, Adjacent protomers of Cx36 (left) and Cx26 (middle) are shown in ribbon representation. Gly227 and Arg233 in Cx36, Arg165 and Ala 171 in Cx26 are shown as sticks. Superposition of Cx36 and Cx26 using Cα atoms of the protomers on the left as reference is shown (right), with Arg233 of Cx36 and Arg165 of Cx26 shown as sticks. **b**, Sequence alignment of multiple connexins near the stereo complementation site. Note that Cx46 and Cx50 are sheep orthologs, whereas the rest are human genes. The red box highlights bulky residues, while the yellow box represents residues with short side chains.

Next, we examined this site in the sequence alignment of a panel of connexins (Fig. 3b). Although the “R” and “short” positions are poorly conserved, almost all connexins harbor an arginine at either position, and fall into either “short-R” or “R-short” category, consistent with our hypothesis. Of known heteromeric gap junction channels, Cx46 and Cx50 in lens fiber cells, both belong to the “short-R” class with short residues being prolines; another example is Cx26 and Cx30, which co-assemble in cochlear supporting cells, are a typical “R-short” and an unusual “Trp-Ile”, respectively. Both examples are consistent with our hypothesis. We further noticed that most α-, γ- and δ-subfamily members are “short-R” type, whereas most β-subfamily members are the opposite “R-short” type. We thereby propose that heteromeric hemichannels might form among α-, γ- and δ-subfamily members, and by members within the β-subfamily (Fig. 4). It is certain that direct biophysical or biochemical investigations are still needed to confirm this proposed specificity of heteromeric hemichannel assembly.

**Fig. 4.**
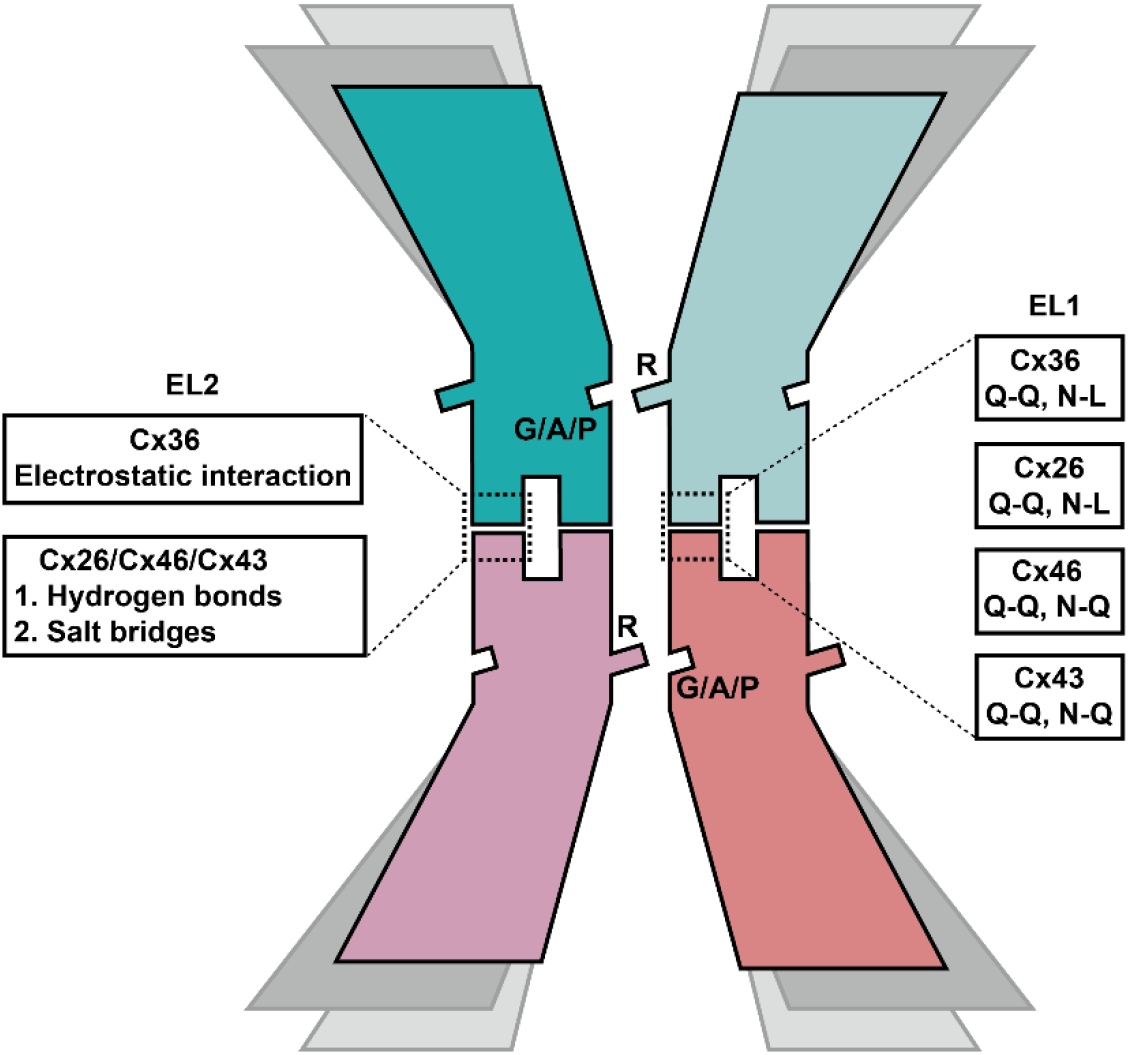
Schematic model of connexin assembly mechanism. The four protomers located in front are colored in teal, pale cyan, light pink and salmon. The four protomers located in the middle and at the back are colored in gray and light gray, respectively. E1-E1 and E2-E2 docking interfaces are labeled with types of interaction and key residues involved. The protrusions and concaves on each protomer represent bulky residues and small side chain residues, respectively.

## Discussion

Connexins are a family of membrane proteins with conserved amino acid sequences and overall structural similarity; yet their functional properties are highly diverse, thus raising a mechanistic hypothesis – the shared connexin architecture is built by conserved amino acids, whereas critical regions are variable to render diverse properties. Of all gap junction channel-forming connexins, the main chain of the channel pore constriction region is structurally conserved, consistent with similar pore profile measured by HOLE program. Their channel properties including ion selectivity and conductance can be very different, likely due to some of poorly conserved pore-lining residues. With determined Cx36 structure, three negatively charged residues from each protomer were mapped at the channel inner surface in close proximity to the pore axis. Related by D6 symmetry, these aspartate and glutamate probably facilitate the permeation of monovalent cations while preventing entrance of anions electrostatically, suggesting a structural basis for the strong cation selectivity of Cx36, analogous to that in members of pentameric Cys-loop receptor family, acetylcholine receptors [60] and γ-aminobutyric acid (GABA) receptors [61], which are cation- and anion-selective, respectively. It has been well established that their opposite ion-selectivity is largely attributed to the opposite charge properties of the channel inner surface. It is thus conceivable that the cation-selectivity of Cx36 shares a comparable mechanism. Consistently, only one of the three pore-lining acidic residues of Cx36 is conserved among all connexins, whereas the other two are often replaced by basic or neutral residues in connexins with little or obscure ion selectivity.

On the other hand, the magnitude of unitary conductance ranges from 15 to 300 pS among connexins, likely independent of ion selectivity [62]. Despite the availability of a panel of connexin structures, yet it remains unclear what the structural determinants account for this widely ranged unitary channel conductance, for which the largely conserved architecture with domestic structural variations elucidated to date is probably insufficient to explain. The structurally disordered intracellular loop or the C-terminal region might play a role. In a single-particle cryo-EM study of another large-pore channel pannexin, structurally disordered C-terminal tail blocked the central entrance, restricting the channel conductance. Caspase cleavage or genetic truncation of the C-terminal tail drastically enlarges the channel conductance by releasing the blockage, allowing the passage of molecules as large as ATP [63]. In comparison, the C-terminal region of Cx36 has been shown to participate in channel gating regulations in responsible to intracellular environments [64]. We speculate that the disordered C-terminal region may form the narrowest restriction along ion-permeation pathway.

The dodecameric assembly of connexin gap junction channel has been a well-acknowledged consensus. In our single-particle analysis, classification of Cx36 projection images revealed evenly distributed projection angles without detectable orientation preferences. Whereas symmetry is difficult to decern from “side” views in parallel to the membrane, six-fold related “top-down” views are readily recognizable, resulted from averaging of the majority of the particles. Interestingly, we also observed putative seven-fold related “top-down” classes containing only a minority of the total particles, consistent with findings described in recently reported analysis of Cx36 similar with this study. Although both studies agreed that these putative seven-fold related Cx36 particles were probably artificial due to detergent extraction, a fundamental question regarding the mechanisms of large-pore channel stoichiometry was raised. Evolutionarily related connexins and pannexins are six- and seven-fold related about their central axis, respectively; another related volume regulatory anion channel (VRAC) LRRC8 can form either homomeric hexamers (LRRC8A or LRRC8D) or heptamers (a functional LRRC8C-8A chimera) [65]. The heptameric structure is more loosely packed, asymmetrical and heterogenous in conformation. Our effort in isolating the putative seven-fold related Cx36 particles by classification failed to yield any reliable reconstruction, likely because these particles are difficult to be distinguished from six-fold related ones and may suffer from similar heterogeneities as seen with the LRRC8C-8A chimera. If mutagenesis of Cx36 could yield a mutant prone to oligomerize into heptamers or tetradecamer, its structural analysis might shed light on the mechanisms by which large-pore channels assemble and molecular evolution in large-pore channel family.

Connexin is a large gene family consisting of 21 members in human genome. When a given cell type expresses two or more connexins, the question of how they may assemble is raised. Some members within the same families have been reported to co-assemble; however, many possible ways of assembly remain. When two connexins are compatible enough with each other, they may oligomerize in a stochastic fashion. Cx46/50 and Cx26/30 are probably good examples. If not compatible at all, a connexin might exclusively form homomers. Cx36 has not been reported to co-assemble with a different connexin to date. Between these two extreme scenarios, it is unclear whether there are preferred homomeric or heteromeric assemblies between or within hemichannels. As the structural collection of representative members of each connexin sub-family becomes complete, it is possible to decern an assembly mechanism that supports the prediction of gap junction channel composition and assembly at neuron-neuron coupling sites, neuron-glia interfaces, and so on. The predicted gap junction channels can thereafter be verified and characterized by a combination of biophysical and biochemical approaches.

## Author contributions

S.C. initiated the project. W.M. carried out protein purification and cryo-EM studies supervised by S.C.. S.C. built the structural model. S.C. and W.M. analyzed the structural model and prepared the manuscript together.

## Acknowledgements

The work was supported by Shanghai Rising-Star Program (20QA1405500), Shanghai Key Laboratory of Translational Medicine on Ear and Nose Diseases (14DZ2260300) and the startup funding provided by Shanghai Insititute of Precision Medicine, Ninth People’s Hospital, Shanghai Jiao Tong University School of Medicine (XK2019006). We thank the staff of the Electron Microscopy Facility at Shanghai Institute of Precision Medicine for providing technical support during data collection. We thank Prof. Jian Wu for helpful advices in model building.

## Extended data

Extended data figs. 1-4 and Extended Data Table 1

