## Supplemental Figures for "Assembly and Cation-selectivity Mechanisms of Neuronal Gap Junction Channel Connexin 36 Elucidated by Cryo-EM"

### Extended Data Figures

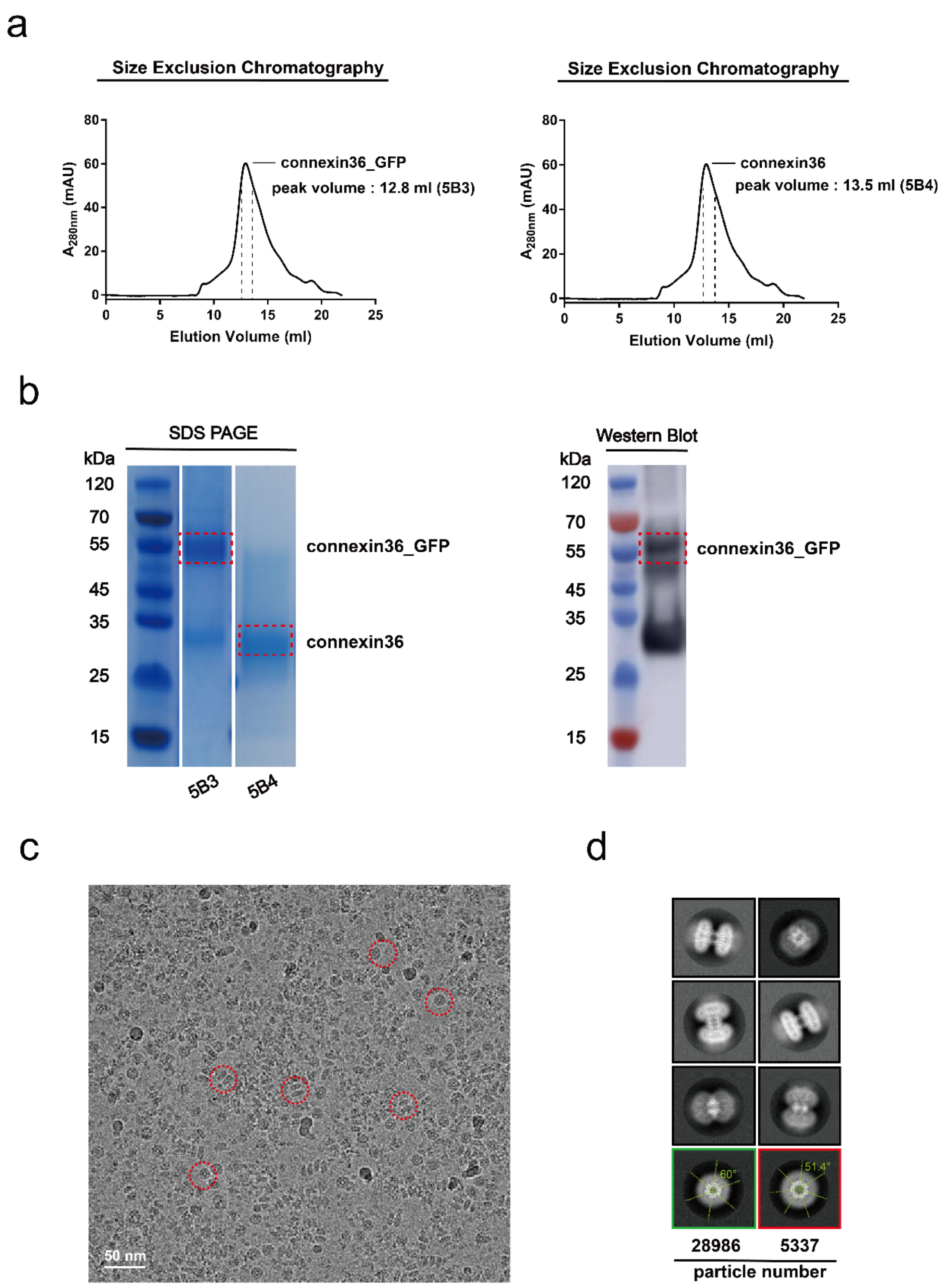

**Extended Data Fig. 1 | Purification and characterization of gap junction protein Cx36. a**, SEC elution profiles of Cx36 in fusion with eGFP (left) and Cx36 free of eGFP (right) in GDN, monitored by UV absorbance. **b**, SDS-PAGE analysis of purified Cx36 in fusion with eGFP and Cx36 free of eGFP (left); western blot analysis of purified Cx36 in fusion with eGFP (right). **c**, A representative cryo-EM micrograph of Cx36. A 50 nm scale bar is shown in the corner. **d**, Representative 2D class averages of Cx36. The green box represents the top-down view of the six-fold symmetric class. The red box represents the top-down view of the seven-fold symmetric class.

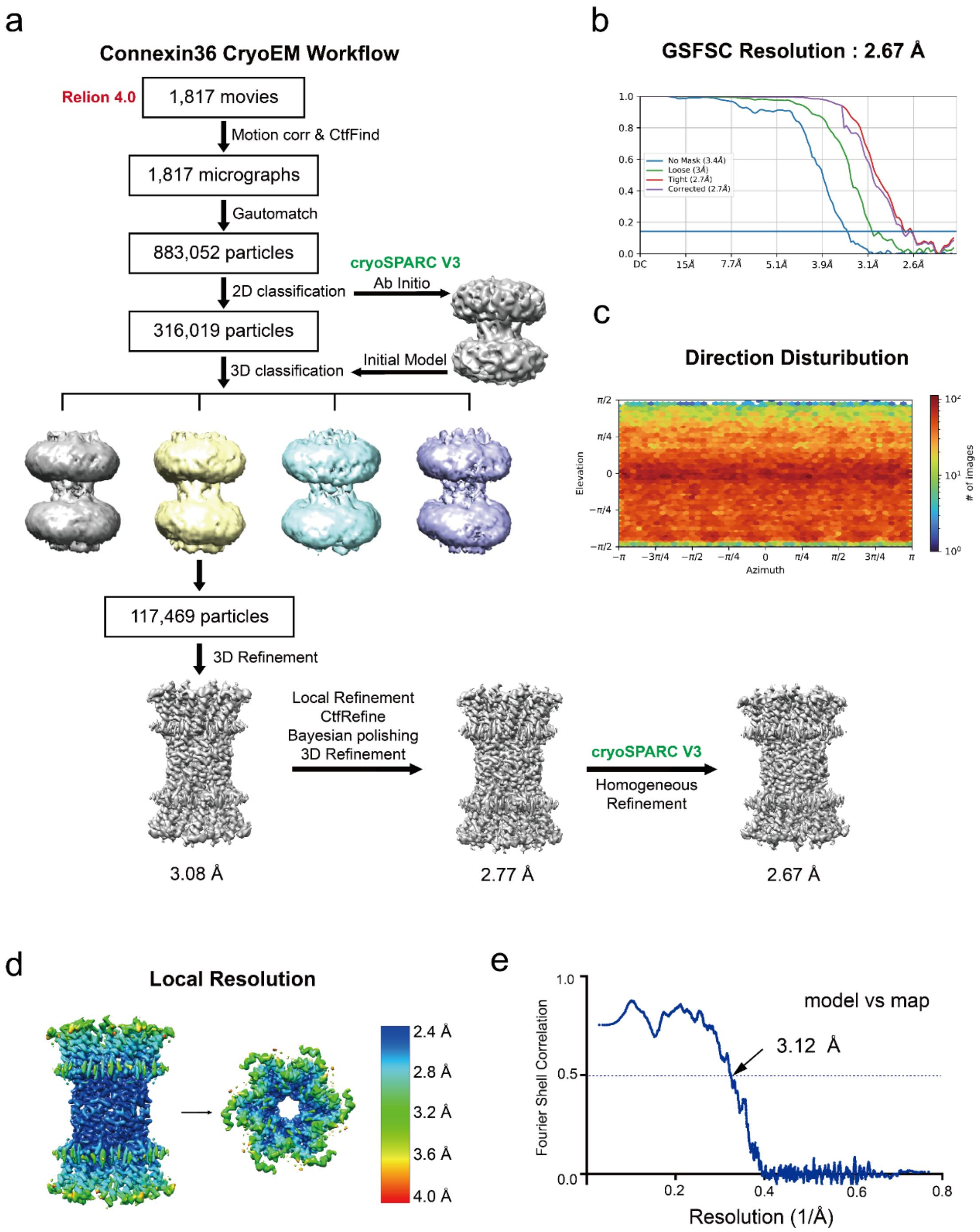

a
Connexin36 CryoEM Workflow
Relion 4.0
1,817 movies
Motion corr & CtfFind
1,817 micrographs
Gautomatch
883,052 particles
cryoSPARC V3
Ab Initio
2D classification
316,019 particles
Initial Model
3D classification
117,469 particles
3D Refinement
Local Refinement
CtfRefine
Bayesian polishing
3D Refinement
cryoSPARC V3
Homogeneous
Refinement
3.08 Å
2.77 Å
2.67 Å
b
GSFSC Resolution : 2.67 Å
No Mask (3.4Å)
Loose (3Å)
Tight (2.7Å)
Corrected (2.7Å)
DC
15Å
7.7Å
5.1Å
3.9Å
3.1Å
2.6Å
c
Direction Disturibution
Elevation
Azimuth
### of images
d
Local Resolution
2.4 Å
2.8 Å
3.2 Å
3.6 Å
4.0 Å
e
model vs map
3.12 Å
Fourier Shell Correlation
Resolution (1/Å)

**Extended Data Fig. 2 | Workflow and validation of cryo-EM data processing. a**, Workflow of cryo-EM data processing for Cx36. **b**, Gold-standard FSC curves for the 3D reconstruction at 2.67 Å with a 0.143 cutoff. **c**, Angular distribution of the particles of the final reconstruction carried out by cryoSPARC homogeneous refinement. **d**, Local resolution estimation by cryoSPARC. **e**, Fourier shell correlation between unmasked cryo-EM map and the refined atomic model indicates an estimated resolution at 3.12 Å with a 0.5 cutoff.

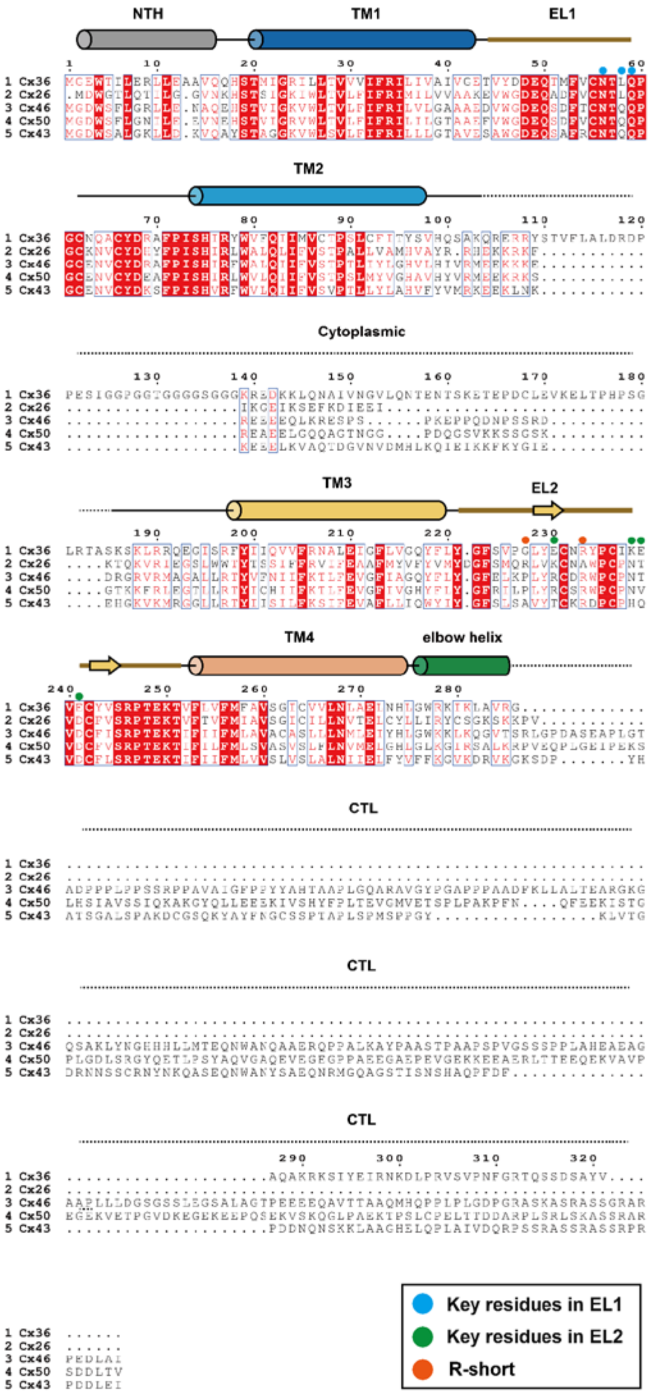
NTH
TM1
EL1
TM2
Cytoplasmic
TM3
EL2
TM4
elbow helix
CTL
CTL
CTL
1 Cx36
2 Cx26
3 Cx46
4 Cx50
5 Cx43
Key residues in EL1
Key residues in EL2
R-short

**Extended Data Fig. 3 | Sequence alignment of five connexins.** Cx36, Cx26, and Cx43 are human sequences, while Cx46 and Cx50 are sheep sequences. All identical residues are boxed in red, similar residues are represented in red font with white background, variable residues are colored in black. Key residues involved in EL1 and EL2 interactions are indicated by colored dots.

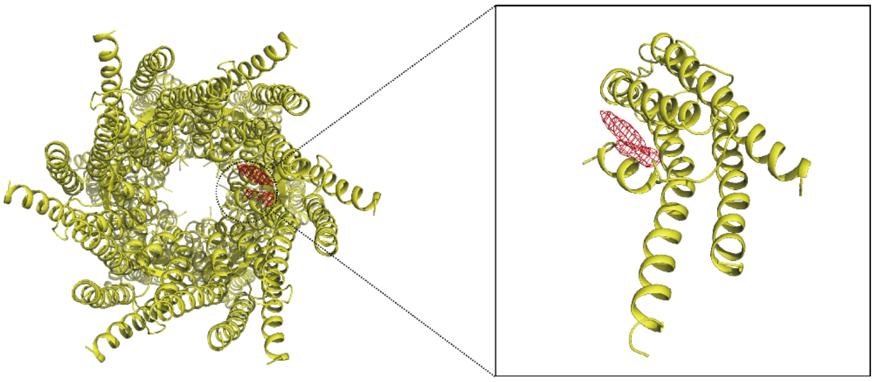

**Extended Data Fig. 4 | Superposition of Cx36 and Cx26.** Align the atomic models of Cx36 and Cx26 (yellow, PDB code 2ZW3) using alpha carbon atoms on the chain A. The atomic model of Cx36 is hidden. The strip density near each protomer in the inner side of the Cx36 channel and the NTH of Cx26 are marked with red dashed circle (left). Magnified view of the strip density in Cx36 and one protomer in Cx26.

**Extended Data Table 1 | Statistics for data collection and structural refinement**

| Statistics for data collection and structural refinement | |
|---|---|
| | Human Connexin 36<br>PDB 8IYG<br>EMDB 35821 |
| **Data collection and processing** | |
| Microscope | FEI Titan Krios |
| Magnification | 81000 |
| Voltage (kV) | 300 |
| Detector | Gatan K3 |
| Electron exposure (e-/Å$^2$) | 50 |
| Defocus range (μm) | 1.5 to 2.5 |
| Pixel size (Å) | 1.1 |
| Symmetry imposed | D6 |
| Initial particle images (no.) | 883052 |
| Final particle images (no.) | 117469 |
| Map resolution (Å) | 2.7 |
| FSC threshold | 0.143 |
| Map resolution range (Å) | 2.4-3.2 |
| **Refinement** | |
| Initial model used (PDB code) | none |
| Model resolution (Å) | 2.7 |
| FSC threshold | 0.5 |
| Map sharpening *B* factors (Å$^2$) | -107.6 |
| Model composition | |
| Non-hydrogen atoms | 1511 |
| Protein residues | 183 |
| *B* factors (Å$^2$) | |
| protein | 145.75 |
| Ligand | 165.37 |
| R.m.s. deviations | |
| Bond lengths (Å) | 0.125 |
| Bond angles (°) | 0.210 |
| Validation | |
| MolProbity score | 1.27 |
| Clash score | 5.14 |
| Poor rotamers (%) | 0 |
| Ramachandran plot | |
| Favored (%) | 99.04 |
| Allowed (%) | 0.96 |
| Disallowed (%) | 0.00 |
